# *multEvol*: An interactive pipeline for multitrait evolutionary analysis in a phylogenetic framework

**DOI:** 10.1101/2022.06.19.496720

**Authors:** Dipendra Nath Basu, Vaishali Bhaumik, Krushnamegh Kunte

## Abstract

Comparative phylogenetic studies based on 2D image data can reveal macroevolutionary patterns and processes by integrating phylogenetic and ecological analyses. The advent of high-throughput imaging facilities has led to the accumulation of large image datasets; however, large-scale data extraction and analysis using these resources can be labour- and computation-intensive. Moreover, there are few standardized methods to analyse multitrait image-based data in a phylogenetic framework. Several packages in the R statistical environment offer utilities for processing and analysing large-scale image data. Techniques for the comparative phylogenetic analysis of multitrait data have also been introduced in other packages. However, most of these approaches are specialized for discrete purposes, and there is a lack of streamlined and efficient methodological pipelines integrating features across packages. Here, we present *multEvol*, an interactive pipeline for the extraction and phylogenetic analysis of shape and colour pattern data from 2D images in the R statistical computing environment. Using this pipeline, shape outlines and colour pattern data for multiple colours can be extracted, visualised, and compared in a phylogenetic framework. Using various utilities provided in the *phytools, RNiftyReg, Momocs, patternize, convevol*, and *geiger* packages, we introduce an interactive interface that allows user inputs for iterative data extraction. This pipeline provides an efficient method for the bulk processing of images of biological specimens from museum collections. As such, it has potential applications in the large-scale morphometric analysis of museum specimens to reveal macroevolutionary patterns in ecology, evolution and integrative biology.

## INTRODUCTION

Earth’s biodiversity is marked by spectacular morphological evolution and diversification of species. Species diversification typically involves evolution along multiple trait axes, which helps organisms adapt to their natural environments and occupy new niches. Comprehensive macroevolutionary analyses of these traits has long posed a challenge to biologists. Quantifying the degree, direction, and rate of morphological change can help elucidate macro- and micro-evolutionary processes in biological systems. However, comparative studies in ecology are frequently limited by a lack of standardized methodology for data extraction and processing [1,2], especially in the bulk analysis of data within a phylogenetic framework. The extracted data may also need to be summarized or otherwise re-structured for use in phylogenetic and statistical analyses, and the methodologies for these are not readily apparent or easily available.

The extraction of shape and colour pattern data from 2D images—such as images of insect wings or plant leaves—offers a versatile means of data collection using relatively inexpensive resources. Comparative studies on diverse taxa, such as bats [3] and butterflies [4], have used shape analysis to examine the evolutionary history and functional constraints of morphological features. Wing colours and colour patterns are extremely diverse and play important roles in mimicry [5], mate recognition [6] and thermoregulation [7], along with other adaptive functions. However, comparative studies on the evolution of complex traits usually focus on relatively closely related species [3,8,9], which may limit the utility of these methods in comparisons across study systems. In addition, standard methods for phylogenetic analysis are typically designed to address specific questions and are incompatible with image data. Therefore, it is important to develop a systematic and interactive pipeline that extracts preliminary data from images and transforms them into phylogenetically corrected formats that are usable for comparative analysis across distantly related taxa.

Here, we present *multEvol*, a pipeline for data extraction from 2D images and phylogenetic analysis of morphological data (Fig. 1) in the R statistical computing environment [10]. The extraction of shape data mostly depends on the *momocs* package [11], which is a dedicated tool for shape analysis in R. The extraction of colour pattern data primarily uses the *patternize* package [12], which allows pattern extraction at the pixel level using a user-defined RGB threshold. In both cases, we have created functions that allow user interaction, which can help ensure proper alignment, harmonic calibration (for shape), and pixel-level pattern selection (for colour) between multiple samples. The outputs can be transformed using phylogenetic principal component analysis (pPCA, package *phytools* [13]), and the PCs can be used for further statistical or phylogenetic analysis, such as estimating the degree of convergence (package *convevol* [14]) or rate of evolution (package *RRphylo* [15]). We demonstrate the utilization of this pipeline using 2D wing images of butterflies. The pipeline can also be used to analyse any continuous variables—such as morphometric data—that have been extracted using other methods.

**Figure 1:**
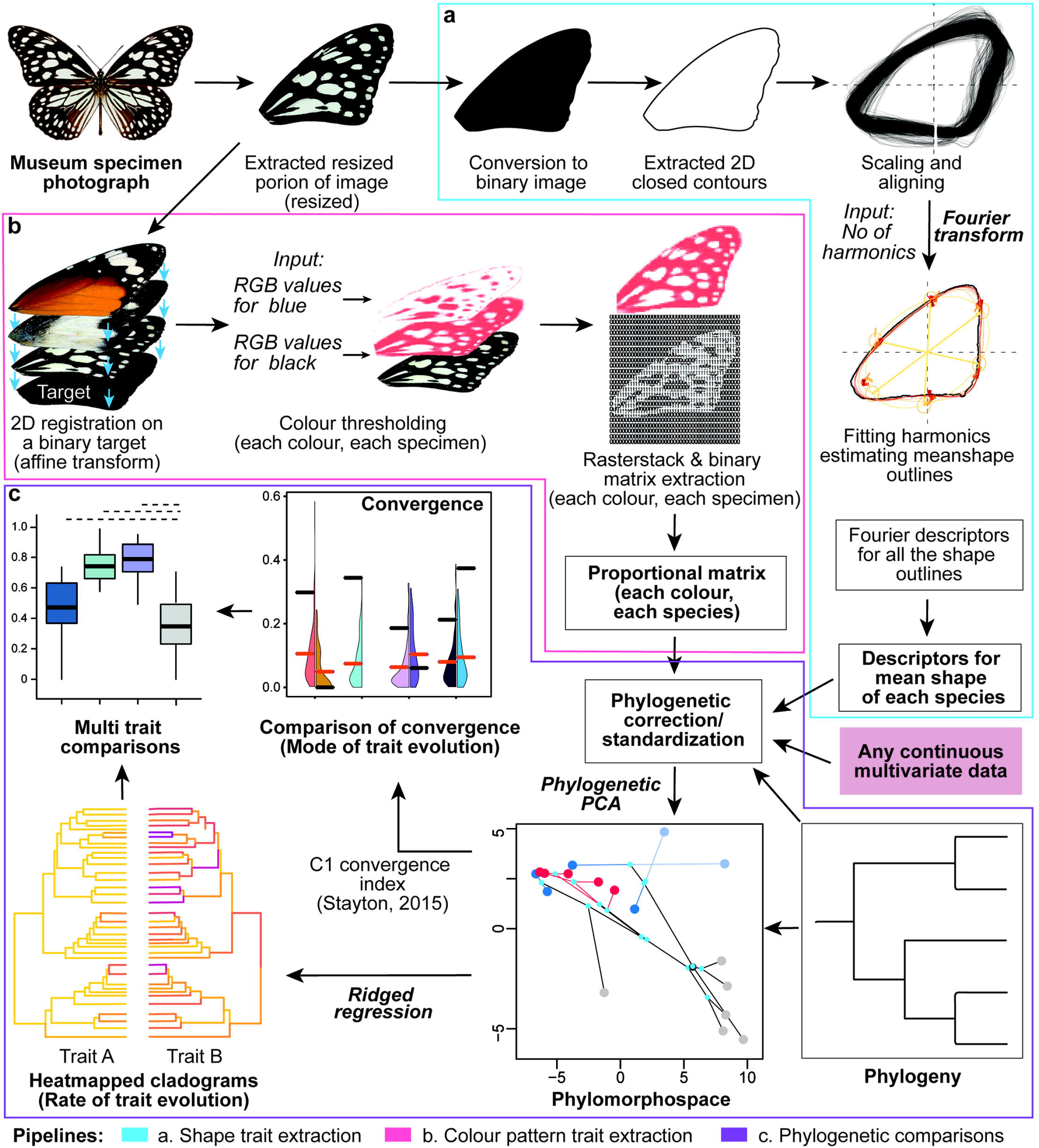
Data extraction, outputs, and potential statistical analyses of shape and colour data using the *multEvol* pipeline. Butterfly wings are photographed and converted into binary images for the extraction of shape (a) or colour pattern (b) data. The 2D shapes are scaled, aligned across all images, and Fourier transformed to obtain the mean descriptors for each species or group. The colour data comprise proportional matrices for multiple colours, where each matrix indicates the pixels that contain the specified colour (0 = none of the images sampled contain the specified colour in that pixel; 1 = all images sampled contain the specified colour in that pixel). The matrices are linearized and used for further analysis, which may include a phylogenetic principal component analysis (PCA) and estimation of convergence or rate of evolution.

## METHODS

### Data extraction

#### Outline shape

In geometric morphometric analyses, biological shapes are estimated through two major approaches: landmark-based analysis and outline analysis. Although both methods retain the relative geometric information of the shapes [16,17], coordinate homology is an essential prerequisite in the landmark method [18]. However, homologous landmarks may not be available in all biological samples, which restricts the utilization of this approach. Additionally, landmarking on the homologous points is a time-consuming process. In contrast, elliptical Fourier analysis (EFA) [17,19,20] is a robust, resource-efficient, and time-efficient method of outline analysis that accurately quantifies 2D closed contours. Unlike other Fourier-based methods, EFA can operate on virtually any shape, including ones that lack positions for homologous landmarks [20]. Contemporary studies on geometric morphometrics are increasingly using EFA to estimate 2D biological shapes [21].

In the EFA approach, the x and y coordinates of closed contours are extracted from the binary images. These outlines are smoothened to a small factor to account for biological aberrations, following which they are centred, aligned, and scaled to remove allometric signals (Fig. 2a). The outlines are size-corrected based on the centroid size. Outline coordinates for each specimen are resampled to introduce homology, and a Procrustes fit is performed on these homologous outline coordinates for better alignment. Resetting the first coordinate of each outline further standardizes the contour estimates. A suitable number of harmonic ranks (a series of ellipses) are then estimated based on the meanshape of the aligned contours (Fig. 2b). Each contour is then represented as a series of calibrated elliptical harmonics (Fig. 2c). These ellipses are defined as the sum of one sine and one cosine curve by decomposing a periodic continuous function implementing an extension of the Fourier series (Fig. 2d). Fourier transformation calculates four coefficients for each harmonic, which are eventually normalized based on the first harmonic to remove the effect of size and rotational variations [17]. These normalized elliptical Fourier descriptors (EFDs) are used as shape variables for further analysis. We implemented a user-friendly interactive function ‘cleqEFA()’ based on utilities provided in R package *Momocs* for outline extraction, alignment, and transformation techniques [11,22].

**Figure 2:**
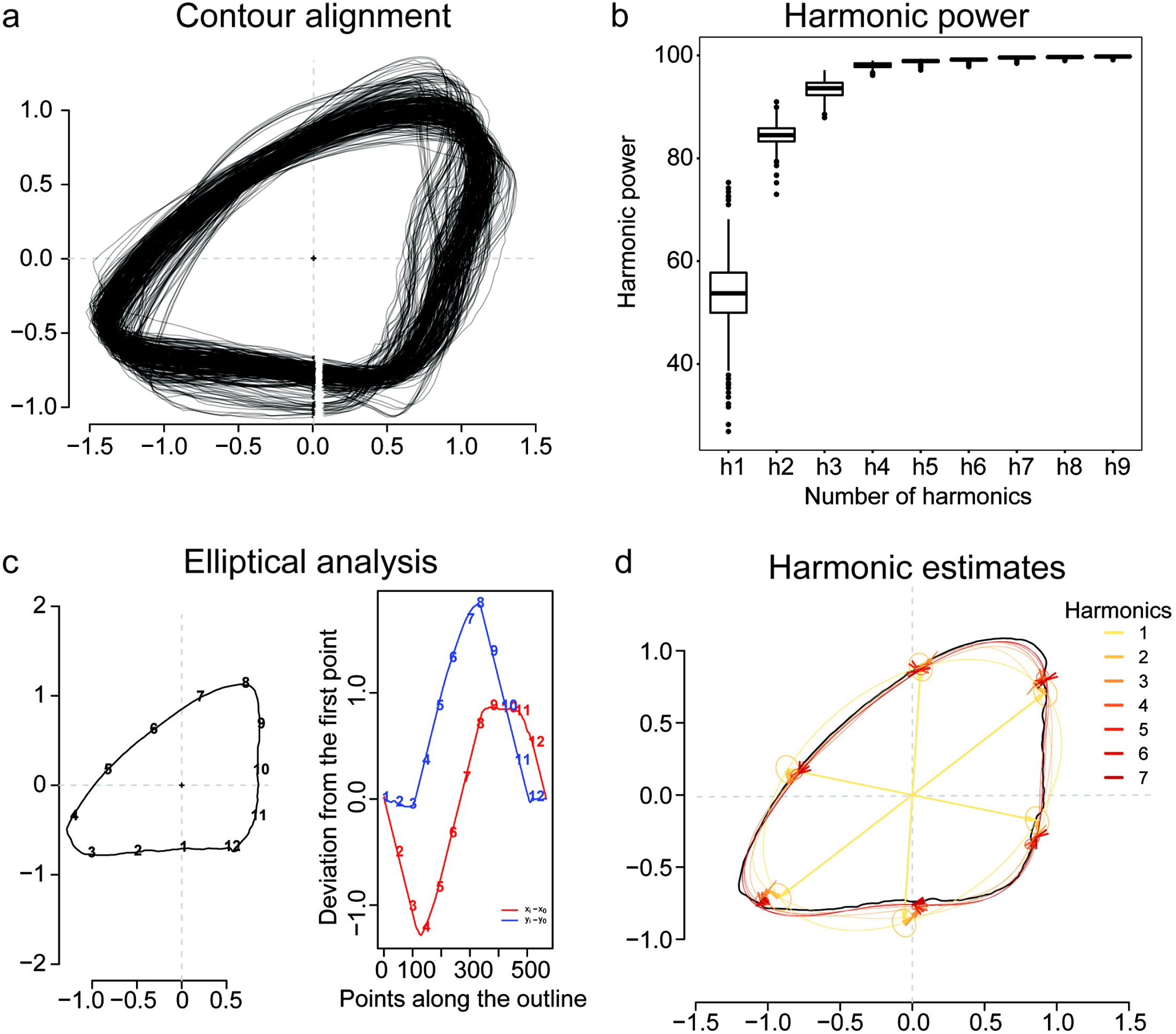
Data processing and extraction of elliptical Fourier descriptors for shape outlines using the ‘cleqEFA()’ function. **(a)** The right forewings (n=265) of butterflies in the *Tirumala* mimicry ring are binarized, and their outline coordinates are extracted. These outlines are scaled and aligned, and the start points of the contours are reset to obtain the final stack of outlines. **(b)** Harmonic power plateaus after the seventh harmonic. **(c)** The harmonic ranks are calibrated to obtain an accurate representation of the outlines. The coordinates of 12 randomly selected equidistant points on the wing contours are plotted directionally to observe the periodic decomposition of the sine and cosine curves from an outline. **(d)** The harmonic sum of a series of fitted ellipses (n=7) on the wing outline—where the higher order ellipse follows the periphery of the lower order ellipse—is Fourier transformed to obtain elliptical Fourier descriptors.

#### Colour pattern

Colour pattern is a complex trait with multiple quantifiable dimensions. Colour pattern data can be quantified in various ways, including relative linear measurements or areas occupied by a particular colour range [23,24] or estimating the shape variance of homologous colour patches across multiple surface [25]. These linear and geometric data can be stored in a presence-absence matrix for each range of colour values (RGB values) on appropriately aligned surfaces. Therefore, this method of pattern extraction makes the data comparable across variable and large image datasets, and can be used for further analysis. The accuracy of these data depends on two components: an efficient image registration approach to introduce homology between samples, and appropriate colour thresholding to extract the spatial and geometric information of homologous patches.

Images can be superimposed to obtain homology through two main approaches: image registration and landmark-based homology. Biological specimens often lack homologous anatomical positions, which restricts the use of landmark-based approaches. Therefore, to obtain homology in a computationally efficient manner, we have opted for a two-step image transformation. In the first step, a binary image template is prepared based on either the meanshape of the 2D surfaces containing colour patterns, or any surface with close resemblance to the meanshape. Subsequently, one specimen of each taxon or species is affine-transformed onto this binary template to obtain a taxon- or species-specific template. In the second step, these templates are used to register all the images using an image registration approach based on similarity of colour intensity (Fig. 1b). The affine transformations are performed iteratively approach using the utilities of the R package *RNiftyReg* [26].

Following image registration, the colour patches that are homologous across multiple surfaces are isolated through thresholding. Most unsupervised thresholding approaches are limited to systems with binary or conspicuous colour patterns (that is, systems in which the surfaces have distinct background and foreground colours), and are not very efficient in systems with multiple colours or continuous colour boundaries. This pipeline uses a supervised RGB thresholding approach where colour boundaries are defined by iteratively sampling from a representative specimen for each species. This approach efficiently extracts colour patterns as raster layer objects for each selected colour range. These raster layers are then converted to binary presence–absence matrices and used for subsequent analyses. RGB thresholding and raster layer extraction are performed using utilities in the R package *patternize* [12].

### Phylogenetic analysis

#### Phylogenetic PCA and size-correction

Most statistical analyses assume that the data have been collected from independent samples. However, this assumption does not hold during data analysis in a phylogenetic framework. Comparative phylogenetic analyses that involve statistical transformation of continuous morphological variables for the estimation of phylogenetic signal or ancestral state reconstruction may require phylogenetic corrections [27]. Statistical comparisons without preliminary phylogenetic corrections show higher variance and are prone to type I error. Hence, this pipeline includes a phylogenetic PCA and offers the option of size correction against body size, as implemented in the R package *phytools* [13].

#### Mode of evolution

Several analytical methods have been developed to estimate non-neutral trends of trait evolution in a phylogenetic framework [9,28]. Estimating these model parameters can reveal the effects of selection and constraints on trait evolution. The ‘pcModels()’ function uses utilities provided in the R packages *APE* [29], *phytools* [13], and *geiger* [30] to fit a phylogenetic tree with a user-defined number of principal components (PCs) and obtain the best fit model. To estimate the extent of convergence in trait evolution, the *convevol* package is used to calculate four convergence indices (C1–C4) [28], which are Euclidean distance-based estimates of convergence among groups of species on a phylomorphospace. Among the various C metrices that can be calculated, C1 denotes the relative convergence in each trait between a group of species, and is comparable across datasets. The *convevol* package also provides a way to simulate data using Brownian motion model parameters to evaluate the significance of the convergence.

#### Rate estimation

Comparative phylogenetic analyses often require estimates of the rate of change in phenotypes across evolutionary time. The rate of trait evolution depends on ancestral state reconstruction, which can be performed in two ways: 1) ancestral state reconstruction following a macroevolutionary model (single- or variable-rate models) within a well sampled clade or closely related clades; and 2) regression-based estimation. A regression-based method is computationally more efficient than Bayesian approaches and can accurately predict the ancestral characters of species belonging to distant clades. The *multEvol* pipeline implements the ridge regression method in the *RRphylo* package [15] to estimate the rate of trait evolution of multiple traits using PCs obtained from the preliminary analysis (Fig. 1c).

## OUTPUT

The main *multEvol* functions for shape extraction take images as input; extract, scale, and align the outlines; and perform elliptical Fourier analysis on those shapes. The output of the ‘cleqEFA()’ function is a list containing the EFDs as a ‘coe’ object, as well as the estimated mean outline for each species as a ‘coo’ object (*Momocs*). The ‘shpComp()’ function extracts the deformation isolines among the meanshapes of different species for comparative visualization. The ‘colData()’ function for colour pattern extraction performs RGB thresholding on images using iterative user inputs in an interactive manner to obtain binary matrices as outputs from each raster object (*patternize*). These matrices are eventually transformed into a proportional matrix for each colour range for each species. The ‘colData()’ function also stores the raster layer objects for each colour range in .png format as confirmation for efficient registration and thresholding, as well as a list of the user-selected RGB value and threshold for each sample. The ‘concat()’ function concatenates the linearized proportional matrices of different colour ranges. The ‘colComp()’ function sums the RasterStacks to visualize the heatmaps of intra- and inter-species colour pattern variation in a comparative manner. The EFDs from outlines, species-level proportional colour pattern data, and any phylogenetically size-corrected continuous multivariate datasets can be subjected to phylogenetic PCA using a phylogenetic tree. The results of phylogenetic PCA are used to visualise inter-species variations in trait space, and the PC scores are stored for further analysis. The ‘dataPrep()’ function fits six macroevolutionary models (Brownian motion [BM], early burst [EB], Ornstein-Uhlenbeck [OU], lambda, kappa, and delta) to the PCs of each trait and stores the AICc values for each model. The ‘cIndex()’ function estimates and stores convergence metrics for each user-defined group of species (regimes) and calculates their significance based on simulations using a BM model. The rate of trait evolution is estimated by the ‘rateEvol()’ function based on the user-defined number of PCs and a phylogenetic tree. This function stores the evolutionary rate for each branch of the phylogeny in a data frame, which can be used to plot a heatmap cladogram.

## EXAMPLE

### Study system

In biological systems, mimicry occurs when a palatable or relatively undefended prey species evolves resemblance to a well-defended (aposematic) species, experiencing reduced predation pressure and deriving a fitness benefit. Aposematic species that resemble each other in a mutualistic interaction are known as Müllerian co-mimics, whereas palatable species that mimic a toxic model in a parasitic interaction are known as Batesian mimics. In this example, we define a mimicry ring as a community that contains at least one well defended aposematic species (Batesian model) and at least one palatable species (Batesian mimic) that resembles the model [31].

Butterflies in a mimicry ring show varying degrees of phenotypic convergence in multiple traits, including colour [32] and colour pattern signals [5,33] as well as locomotor cues [34]. These traits may evolve under different models of selection and at varying rates within and between species [4,23]. The *Tirumala* mimicry ring endemic to the Western Ghats biodiversity hotspot in India consists of five unpalatable models with a shared phenotype and shared ancestry, and three distantly related palatable species (Batesian mimics) that have evolved to resemble the toxic models (Fig. 3) [31,35]. The mimics comprise two species of female-limited mimics (*Pareronia hippia* and *P. ceylanica*), where the males are non-mimetic, and one species of sexually monomorphic mimic (*Papilio clytia* f. *dissimilis*) where both sexes share the mimetic phenotype. Here, we examine the evolution of wing shape and colour patterns in this mimicry ring using the *multEvol* pipeline to demonstrate its utility in investigating the evolution of multiple traits under various degrees of mimetic selection. For phylogenetic analyses, we first prepared a time-calibrated phylogeny consisting of the models, mimics, and some of their respective sister species. We then extracted wing shape and colour data from the forewing images of all species and estimated the degree of convergence between models and mimics, as well as the rate of evolution of the two traits, as described above.

**Figure 3:**
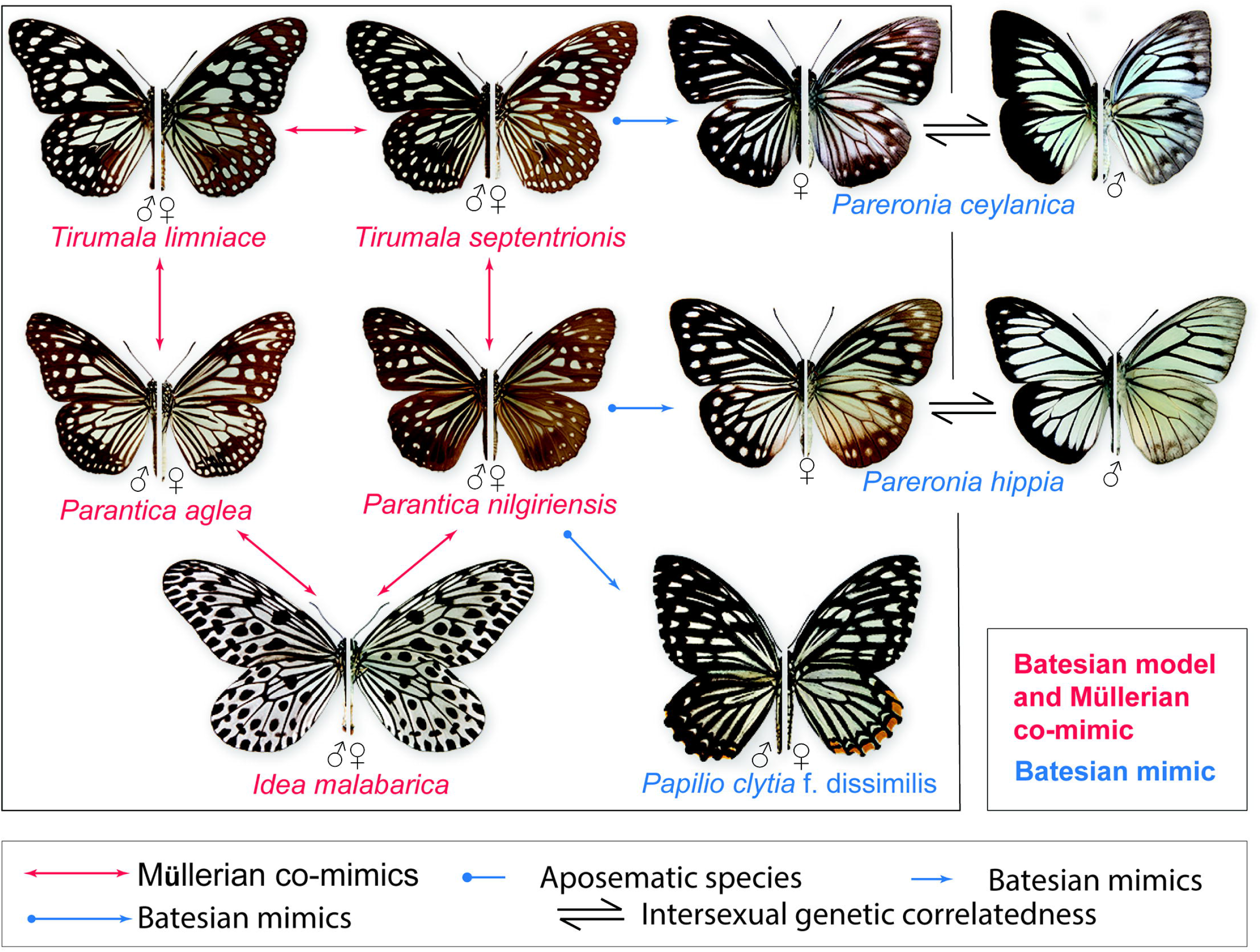
Müllerian co-mimics and Batesian mimics in the *Tirumala* mimicry ring of the Western Ghats, India. The five Müllerian co-mimics (red arrows) have shared ancestry and act as Batesian models for the Batesian mimics (blue arrows), which are distantly related. *Pareronia* spp. are female-limited mimics, where the females mimic the models and the males are non-mimetic.

### Data extraction and preliminary analysis

Using digital SLR cameras and lenses, we photographed one forewing (dorsal surface) of each specimen and extracted the wing shape and colour pattern data. Images were captured under a uniform camera flash units because variation in colour tones under different light conditions may produce different results. We extracted shape data from 265 images using the ‘cleqEFA()’ function, and colour data from 200 images using the ‘colData()’ function. Seven harmonic ranks were calibrated to explain the outlines with 99% accuracy (Fig. 2b). The EFDs from these seven harmonics were normalized against those of the first harmonics, resulting in six remaining EFDs. Data were extracted for three colour ranges— melanistic (including black and brown), red/orange, and blue/green—with a resampling factor of three, and the resulting proportional matrices for each colour range were linearized and concatenated for each species. The EFD datasets and concatenated colour pattern data were used for a phylogenetic PCA, and six evolutionary models were fit to the resulting PCs.

## RESULTS

The first two PCs explained 96.23% and 98.1% of the variance in shape and colour pattern data, respectively. Of the six models compared, the lambda model of evolution had the highest support for the first two of shape (AICc values: PC1, −83.21; PC2, −110.51) and color (AICc values: PC1, 228.76; PC2, 174.03). The meanshapes of each model and mimic species were reconstructed from their respective EFDs. For each model–mimic pair, the shape changes were visualized as deformation isolines between the meanshapes of each species (Fig. 4a). In contrast to sexually monomorphic mimics and mimetic females of female-limited mimics, the non-mimetic males showed multiple local deformations (red areas in Fig. 4a). The summed raster layers were plotted as heatmaps to visualise intra-species colour variation (that is, variation in colour patterns between samples of a single species; plots along the top and left margins in Fig. 5a). Pairwise differences in the species-level proportional raster layers were plotted as heatmaps to indicate inter-species differences in colour patterns between each model–mimic pair (Fig. 5a). Similar to the trend observed in wing shapes, non-mimetic males showed more differences in colour patches compared to models, whereas sexually monomorphic and female-limited mimics were similar to models. The phylomorphospaces for shape and colour pattern analysis indicated an overlap between models and mimics (Figs. 4b and 5b). In contrast to mimetic females, the non-mimetic males of female-limited mimetic species were relatively distant from the traitspace occupied by models and mimics. The convergence indices of various species combinations (only Batesian mimics, only aposematics, sexually monomorphic mimics + aposematics, and female-limited mimics + aposematics) were plotted using the shape and colour pattern data to visualize the distribution of C1 values in 100 simulated datasets (Fig. 6a). There was significant convergence in wing shape and colour patterns within models and between models and female-limited mimics, as well as in wing shape between models and sexually monomorphic mimics (star symbols in Fig. 6a). The evolutionary rates of wing shape and colour pattern were plotted onto heatmap cladograms, which showed that lineages containing models or mimics generally evolved faster (Fig. 6b).

**Figure 4:**
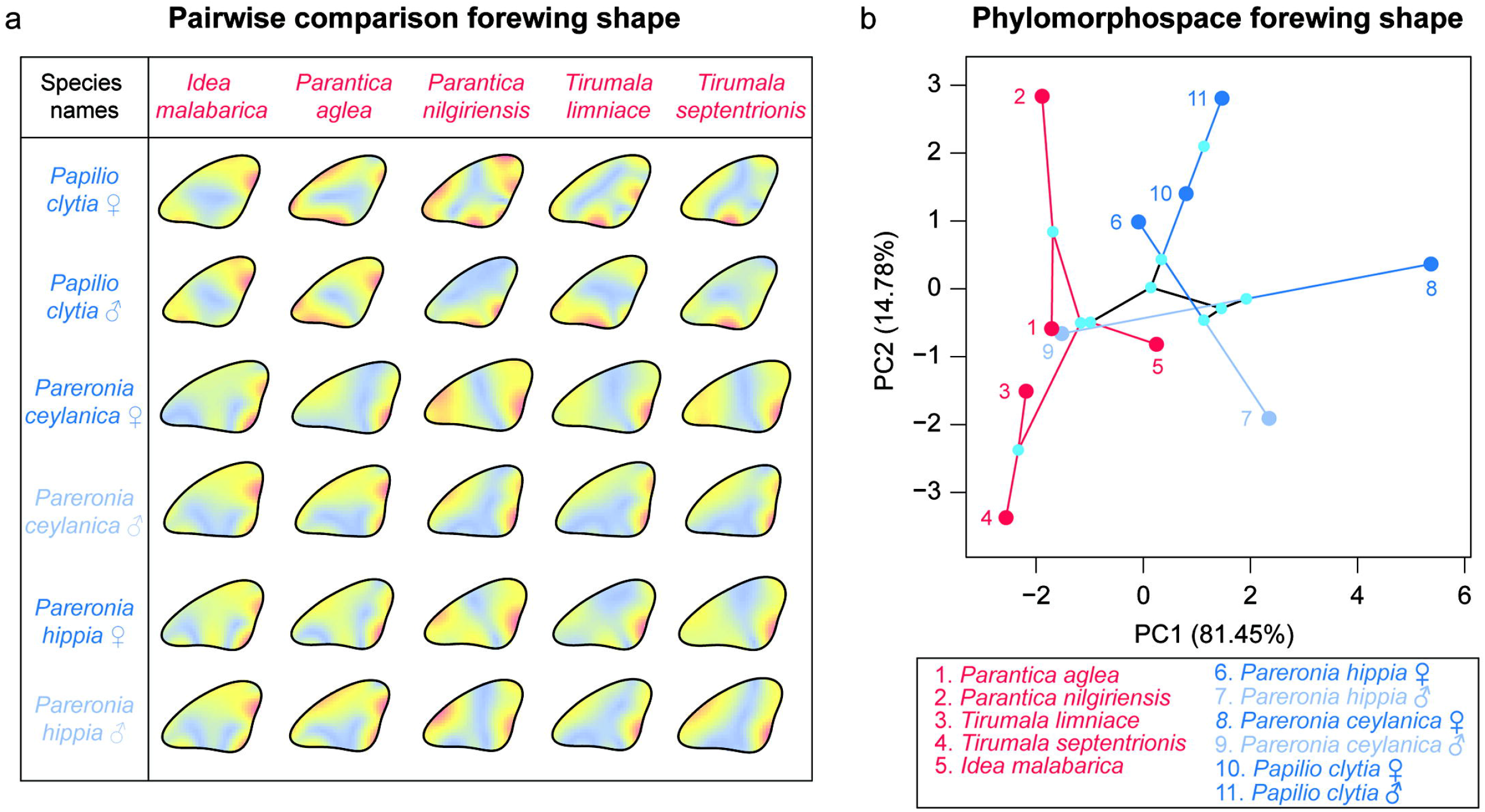
Inter-species variation in wing shape among mimetic butterflies. **(a)** Pairwise comparisons of deformation isolines (scale: blue=low deformation, yellow=moderate deformation, red=high deformation), showing the changes in wing shape in model–mimic species pairs in the *Tirumala* mimicry ring. **(b)** A morphospace plot based on the phylogenetic principal component analysis of shape descriptors, showing the relative positions of aposematics (red), mimics (dark blue), and non-mimetic males of female-limited mimics (light blue) in the traitspace.

**Figure 5:**
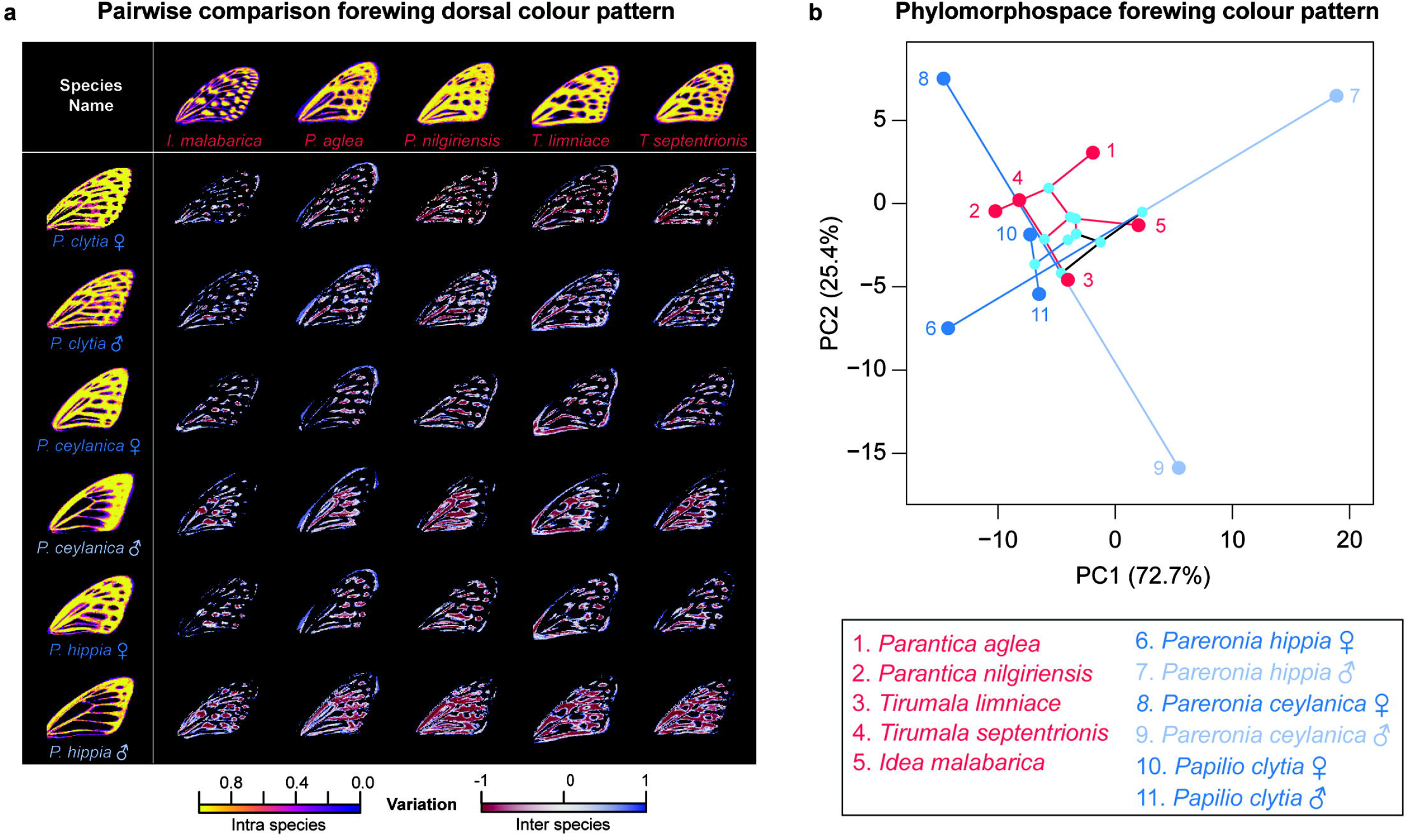
Intra- and inter-species variation in wing colour patterns among mimetic butterflies. **(a)** Heatmaps along the top and left margins show intra-species variations in colour patterns, and the remaining heatmaps exhibit inter-species differences in colour patterns in model– mimic species pairs in the *Tirumala* mimicry ring. The heatmap scale is shown at the bottom right. **(b)** A morphospace plot based on the phylogenetic PCA of colour pattern data, showing the relative positions of aposematics (red), mimics (dark blue), and non-mimetic males of female-limited mimics (light blue) in the traitspace.

**Figure 6:**
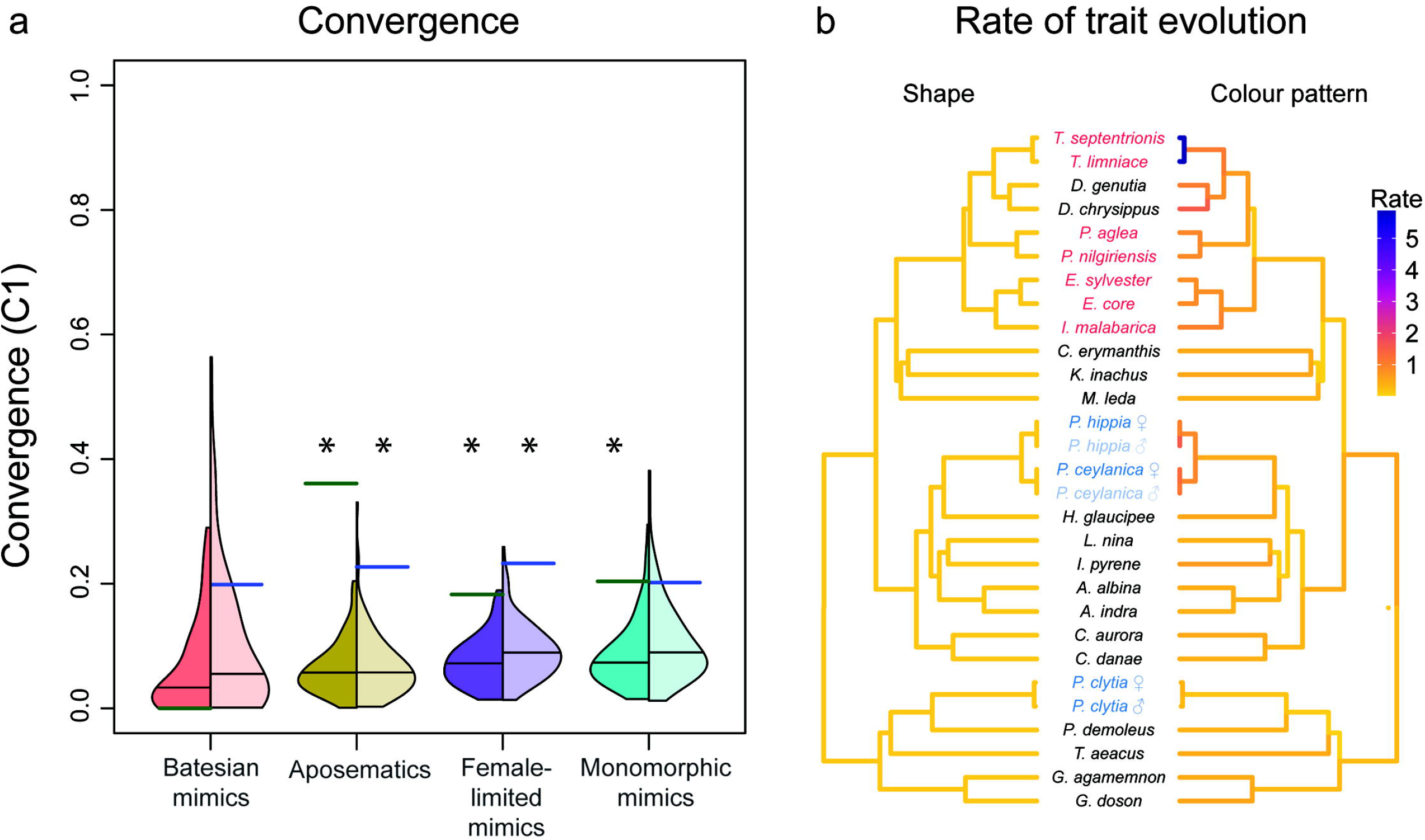
Evolutionary characteristics of forewing shape and forewing dorsal colour patterns in the *Tirumala* mimicry ring. **(a)** Convergence among Batesian mimics (distantly related palatable species; includes female-limited dimorphic and sexually monomorphic mimics); among aposematics (toxic species with similar colour patterns); between aposematics and female-limited dimorphic mimics; and between aposematics and sexually monomorphic mimics. C1 values (red: actual C1; black: mean C1 across 100 simulations) are shown for forewing shape (left) and forewing dorsal colour patterns (right). A star indicates significant convergence relative to the convergence estimates from 100 simulations. **(b)** Rates of evolution of forewing shape and forewing dorsal colour patterns. Colour coding for species names indicates aposematics (red), Batesian mimics (dark blue), and the males of female-limited Batesian mimics (light blue).

## CONCLUSION AND RECOMMENDATIONS

The *multEvol* pipeline can be used to compare the distribution of colour ranges across species, and the interactive features allow the iterative selection of appropriate harmonics, RGB values, and colour pattern matrices across multiple samples. This is especially useful for large-scale supervised data collection from museum collections or field-collected specimens. Museum collections worldwide represent a vast resource of historical DNA; however, preserved specimens have limited applicability as sources of biological tissue. As such, the systematic photographing and cataloguing of museum specimens can improve their long-term usefulness as sources of morphological data [4,36]. The utilization of museum specimens alleviates several taxonomical and logistical limitations [37,38] and enables the testing of various macroevolutionary hypotheses. The pipeline presented here provides the potential for such large-scale comparative analyses, and we hope that the methods described and pipeline presented here will prove useful to researchers studying shape and colour pattern variation in biological systems.

## Author Contributions

DNB and VB conceptualised the analytical pipeline, developed the code, and conceptualised and performed phylogenetic analyses. KK conceived the broad research questions, supervised the research, and contributed specimens for image analysis. DNB and VB wrote the manuscript with inputs from KK.

## Data availability

The functions and their descriptions are available on GitHub (https://github.com/DipendraBasu/multEvol-0.1).

